# Acute alcohol and chronic drinking bidirectionally regulate the excitability of prefrontal cortex vasoactive intestinal peptide interneurons

**DOI:** 10.1101/2023.03.07.531614

**Authors:** Shannon M. Thompson, Anthony S. Ferranti, Max E. Joffe

## Abstract

The prefrontal cortex (PFC) regulates drinking behaviors and affective changes following chronic alcohol use. PFC activity is dynamically modulated by local inhibitory interneurons (INs), which can be divided into non-overlapping groups with distinct functional roles. Within deeper layers of neocortex, INs that express either parvalbumin or somatostatin directly inhibit pyramidal cells. By contrast, the plurality of all remaining INs express vasoactive intestinal peptide (VIP), reside within superficial layers, and preferentially target other types of INs. While recent studies have described adaptations to PFC parvalbumin-INs and somatostatin-INs in alcohol use models, whether ethanol or drinking affect the physiology of PFC VIP-INs has not been reported. To address this gap, we used genetically engineered female and male mice to target VIP-INs in layers 1-3 of prelimbic PFC for whole-cell patch-clamp electrophysiology. We found that ethanol (20 mM, ∼0.09 BEC) application to PFC brain slices enhances VIP-IN excitability. We next examined effects following chronic drinking by providing mice with 4 weeks of intermittent access (IA) ethanol two-bottle choice in the home cage. In these studies, VIP-INs from female and male IA ethanol mice displayed reduced excitability relative to cells from water-only controls. Finally, we assessed whether these effects continue into abstinence. After 7-11 days without ethanol, the hypo-excitability of VIP-INs from male IA ethanol mice persisted, whereas cells from female IA ethanol mice were not different from their controls. Together, these findings illustrate that acute ethanol enhances VIP-IN excitability and suggest these cells undergo pronounced homeostatic changes following long-term drinking.

## 1 Introduction

Breakthrough treatments are needed to meet the ongoing public health challenges related to maladaptive drinking and alcohol use disorder (AUD) (Grant et al., 2017; Whiteford et al., 2013). Developing new neuropharmacological treatments necessitates a deeper understanding of the mechanisms through which ethanol alters brain circuit function. Studies in humans and in animal models suggest that difficulty in moderating drinking and affective disruptions during abstinence are each associated with the prefrontal cortex (PFC) (Blaine et al., 2020; Koob, 2014). In addition, differences in PFC function, at the population level, have been related to heavy drinking or AUD diagnosis (Medina et al., 2008; Sorg et al., 2012; Wang et al., 2016). A better understanding of how specific PFC cell types are affected by ethanol and drinking should inform new avenues for treatment development.

PFC pyramidal cells regulate drinking through long-range glutamatergic projections to subcortical structures (Klenowski, 2018; Seif et al., 2013; Siciliano et al., 2019). Nonetheless, local inhibitory interneurons (INs) orchestrate the activity of pyramidal cells. At the highest level, neocortical interneurons can be subclassified by mutually exclusive expression of parvalbumin (PV), somatostatin (SST), or (in rodents) the 5HT_3_ serotonin receptor (Lee et al., 2010; Rudy et al., 2011). Accumulating evidence indicates that acute ethanol, voluntary drinking, and dependence all alter the activity of PFC PV-INs and SST-INs (Dao et al., 2021; Dao et al., 2020; Ferranti et al., 2022; Fish and Joffe, 2022; Hughes et al., 2020; Joffe et al., 2020b; Li et al., 2021; Trantham-Davidson et al., 2014; Trantham-Davidson et al., 2017). Considering these two IN subtypes regulate PFC circuit function through distinct subcellular targets and mechanisms, it is unsurprising that prior studies performing head-to-head comparisons each observed contrasting adaptations to PV-INs and SST-INs in alcohol use models (Hughes et al., 2020; Joffe et al., 2020b). In addition, other specialized IN subtypes most likely undergo distinct adaptations in response to ethanol. At this point, however, very little is known regarding how ethanol affects the physiology or function of any subclass of 5HT_3_-IN.

Vasoactive intestinal peptide (VIP) is exclusively expressed by GABAergic neurons in neocortex (Pronneke et al., 2015; Tasic et al., 2018). VIP-INs represent the plurality of 5HT_3_-INs and reside primarily within superficial layers of neocortex (Lee et al., 2010), and most VIP-INs are bipolar cells that co-express calretinin (Cauli et al., 2014; Kawaguchi and Kubota, 1997). VIP-INs display extremely high membrane resistance (R_m_), rendering them exquisitely responsive to stimulation (Anastasiades et al., 2018; Lee et al., 2013; Pronneke et al., 2015; Schuman et al., 2019). Unique among IN subtypes, VIP-INs display minimal innervation of pyramidal cells and instead preferentially target SST-INs (Anastasiades et al., 2018; Fu et al., 2015; Lee et al., 2013; Melchitzky and Lewis, 2008; Pfeffer et al., 2013; Pi et al., 2013). Through this disinhibitory motif, VIP-INs are activated by subcortical nuclei to decrease spontaneous SST-IN activity, increase the gain through cortical circuits, and ultimately facilitate pyramidal cell activity (Audette et al., 2018; Lee et al., 2013; Pi et al., 2013; Williams and Holtmaat, 2019). VIP-INs are therefore well-positioned to regulate the effects of drugs and other salient experiences; nonetheless, we are not aware of any published studies that have reported how ethanol or drinking affect the physiology of PFC VIP-INs.

To address this knowledge gap, we aimed to test whether alcohol alters the physiology of PFC VIP-INs. Using targeted whole-cell patch-clamp techniques, we discovered that acute application of a modest concentration of ethanol (20 mM / 0.09 mg/dL) increases the excitability of PFC VIP-INs in mice. Next, to ask whether chronic drinking induces similar or related adaptations to VIP-INs, we provided mice with intermittent access (IA) ethanol. In contrast to the acute application studies, we found that VIP-INs from IA ethanol mice displayed decreased spike-firing relative to cells from water-only controls, suggesting that chronic drinking can induce a homeostatic reduction in VIP-IN excitability. Finally, we found that VIP-INs underwent sex-dependent changes during 7-11 days of abstinence: VIP-INs from male IA ethanol mice exhibited persistent hypo-excitability relative to controls, whereas VIP-INs from female IA ethanol mice were not different from respective controls. Together, these studies describe adaptations to PFC VIP-INs following acute ethanol, chronic drinking, and abstinence, suggesting that neocortical disinhibitory microcircuits contribute to acute and chronic adaptations related to alcohol use.

## 2 Material and Methods

### 2.1 Mice

Mice were housed on a standard 12-hour light cycle (on at 6:00 AM). Mice were bred on a congenic C57BL6/J background to express tdTomato in PFC VIP-INs. Female VIP-Cre mice (Madisen et al., 2010) (*Vip*^*tm1(cre)Zjh*^*/J*, Jackson Laboratories, Stock No: 010908) were crossed with male “Ai9” mice (Taniguchi et al., 2011) (B6.Cg-*t(ROSA)26Sor*^*tm9(CAG-tdTomato)Hze*^*/J*, Jackson Laboratories, Stock No: 007909). We classified mice as female or male based only on external genitalia, and all mice were used for experiments. Comparable trends for the acute effects of ethanol on VIP-IN physiology emerged in neurons from both female mice and male mice, so data were pooled for the primary analyses. Given sex differences in voluntary drinking in rodents, all analyses for IA ethanol experiments accounted for sex as a separate variable. For the IA ethanol experiments, mice were group-housed (2-5 per cage) in clear polysulfone individually ventilated cages until IA ethanol experiments began, no earlier than 6-weeks of age, after which they were individually housed. For all other studies, mice were always group-housed.

### 2.2 Electrophysiology

We used whole-cell patch-clamp electrophysiology to assess VIP-IN membrane and synaptic physiology. As in previous studies (Joffe et al., 2022; Joffe et al., 2020a), we rapidly prepared 300-μm coronal brain slices containing prelimbic PFC using an NMDG-based cutting and recovery solution (in mM): 93 NMDG, 2.5 KCl, 0.5 CaCl_2_, 10 MgSO_4_, 1.2 NaH_2_PO_4_, 30 NaHCO_3_, 25 glucose, 20 HEPES, 5 Na-ascorbate, and 3 Na-pyruvate; pH adjusted to 7.3 with HCl, 300-310 mOsm. All solutions were bubbled with 95% O_2_ / 5% CO_2_. Slices recovered in standard artificial cerebrospinal fluid (ACSF) at 20-24 °C for 1-6 hours before recordings. ACSF contained (in mM): 119 NaCl, 2.5 KCl, 2 CaCl_2_, 1 MgCl_2_, 1 NaH_2_PO_4_, 26 NaHCO_3,_ and 11 glucose; 292-296 mOsm. Slices were transferred to a recording chamber perfused with 30-32 °C at 2 mL/min under an upright microscope. We targeted VIP-INs in layers 1-3 prelimbic PFC via red, tdTomato fluorescence and patched cells using pulled glass pipettes (5-7 MΩ) filled with a potassium-based internal solution (in mM): 125 K-gluconate, 5 NaCl, 0.2 EGTA, 10 HEPES, 4 MgATP, 0.3 Na_2_GTP, 10 Tris-phosphocreatine; pH adjusted to 7.3 with KOH, 292-296 mOsm.

Cells were dialyzed with internal solution for 5 minutes before undergoing a series of 1-sec current injections [-30 to +150, Δ5 pA] to assess VIP-IN membrane properties. We examined several physiological properties, including: (1) resting membrane potential (V_m_), (2) membrane resistance (R_m_), (3) hyperpolarization sag, (4) rheobase, and (5) spike firing input-output curve. R_m_ was calculated as the best-fit slope from three hyperpolarizing current injections [-5, -10, -15 pA]. The hyperpolarization sag ratio was quantified as the difference between the peak hyperpolarization and the steady-state hyperpolarization normalized to the difference between the steady-state hyperpolarization and the resting potential. The sag ratio was calculated as the average value across three consecutive current injections [-20, -25, -30 pA]. The rheobase and spike-firing input-output curves were collected during and after depolarizing current injections. Rheobase was determined as the minimum current step that initiated at least one action potential. Action potentials were identified using a threshold search event detector and the number of spikes per current step were reported. Cells that spontaneously fired actional potentials were discarded. After current-clamp recordings, cells were voltage-clamped at -80 mV to electrically isolate spontaneous excitatory postsynaptic currents (sEPSCs) (Joffe et al., 2020b). sEPSCs were identified using a template search event detector.

For ethanol application studies, slices were incubated in 20 mM ethanol for at least one hour before, and throughout, all recordings. Preliminary studies found minimal effect of 5-minute ethanol applications. Control recordings in standard ACSF were always recorded from the same animals on the same recording days in a double-alternating, counterbalanced manner. In some studies, we prepared acute brain slices after 4 weeks of voluntary drinking. Slices were prepared either the morning after a night of ethanol access or 7-11 days following forced abstinence. We recorded from 3-6 cells per animal and did not find evidence for any correlations between an animal’s ethanol consumption and any physiological parameter. The experimenter was blinded to condition for all studies following chronic drinking.

### 2.3 IA ethanol

As in our previous studies (Joffe et al., 2020b, 2021), we modeled high levels of volitional alcohol drinking by an intermittent access (IA) schedule. For three 24-hour periods per week, ethanol was provided in home cages. Ethanol was removed for 24-48 hours between each exposure, and mice had constant access to food and water throughout all experiments. Ethanol was diluted from 95% undenatured ethyl alcohol (Decon Labs, Incorporated). Ethanol and water were delivered by conical tubes capped with rubber stoppers (Fisherbrand 14-135H, Fisher Scientific) and open tip straight tubes (OT100, Ancare). Fluid consumed was calculated by measuring tube weight before and after each session, subtracting the average drip (0.2 g) from all values. Ethanol was generally provided and removed 3-4 hours prior to the transition to the dark cycle. The ethanol concentration increased over the first three sessions [3, 6, 10%] after which 20% ethanol was provided for the remainder of the study.

### 2.4 Statistics

The number of cells and mice are denoted by “n” and “N”, respectively, for each experiment. ClampFit 11.2 (Molecular Devices) was used to for primary analysis and subsequent analyses were performed in Excel (Microsoft) and Prism 9.4 (GraphPad). Data are presented as mean ± standard error or as box plots. Data were analyzed using two-tailed Student’s t-test, or Mann-Whitney test if a dataset failed a Shapiro-Wilk normality test. Two-way ANOVA or mixed-effects analysis with Bonferonni post hoc comparisons were also used to detect significance.

## 3 Results

### 3.1 Minimal sex differences in membrane physiology of PFC VIP-Ins

Within neocortex, VIP is selectively expressed by disinhibitory interneurons enriched in superficial layers **(Figure 1a)**. To target these cells in a high-throughput manner, we bred mice to express red fluorescent protein in PFC VIP-INs by crossing homozygous VIP-Cre female mice (Madisen et al., 2010) with male mice harboring a Cre-dependent tdTomato locus (Taniguchi et al., 2011). We made acute brain slices from adult (8-20-week) mice and performed whole-cell patch-clamp electrophysiology experiments to examine VIP-IN membrane physiology parameters **(Figure 1b)**. As reported in previous studies (Anastasiades et al., 2018; Goff and Goldberg, 2019; Kawaguchi, 1997; Porter et al., 1998; Pronneke et al., 2015), we found PFC VIP-INs to be highly excitable cells with considerable heterogeneity in spike-firing patterns and subthreshold properties.

**Figure 1.**
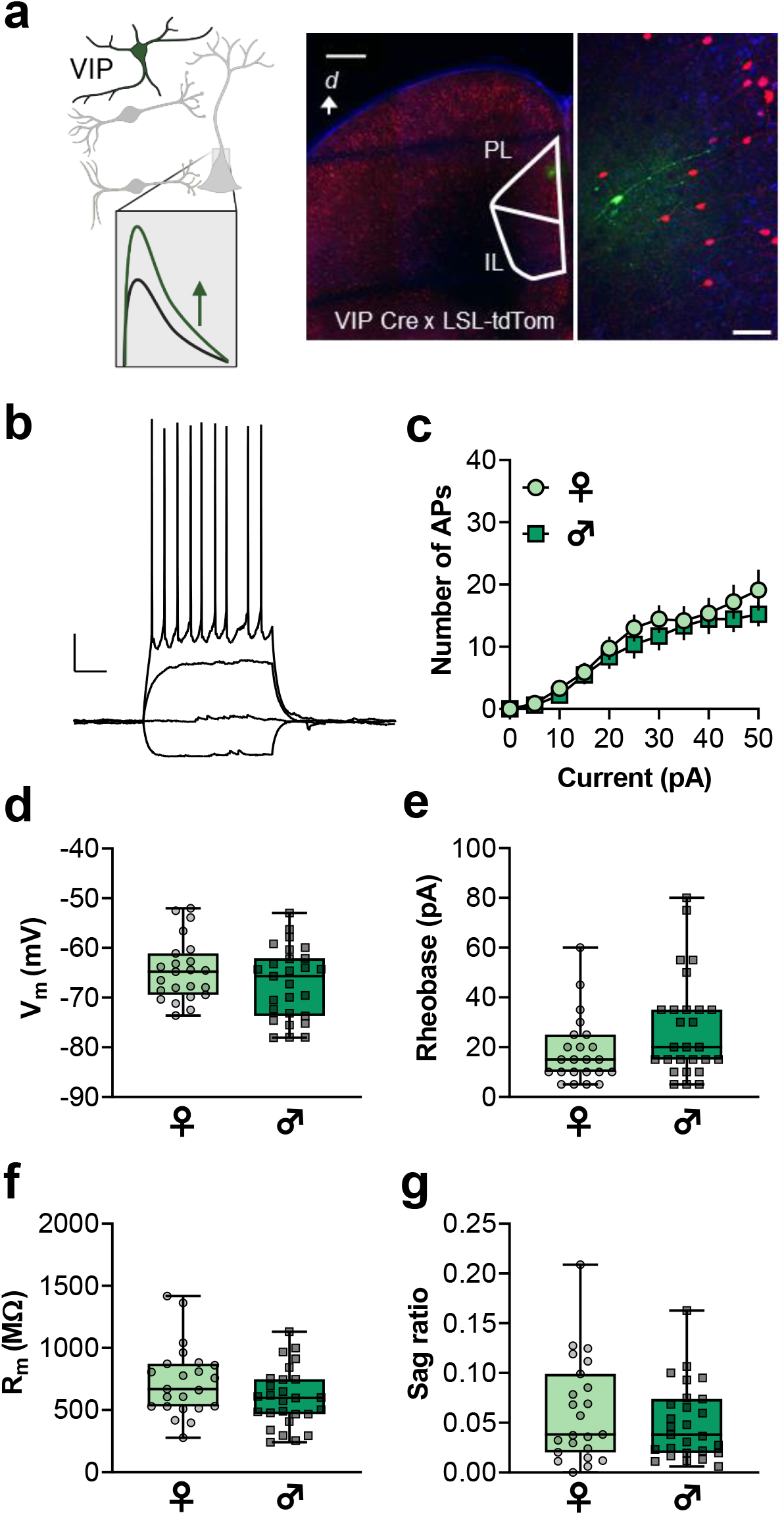
Minimal sex differences in membrane physiology of PFC vasoactive intestinal peptide inhibitory interneurons (VIP-INs). **(a)** VIP-INs inhibit other IN subtypes to enhance excitatory postsynaptic potentials in pyramidal cells. VIP-Cre mice were crossed with Ai9 mice (Rosa26-Lox-STOP-Lox-tdTomato) to generate mice expressing fluorescent proteins in VIP-INs. A representative image of a coronal brain slices depicts a representative cell filled with Alexa Fluor 488 green dye. d, dorsal; IL, infralimbic; PL, prelimbic. Scale bars 500 μm and 50 μm. **(b)** Representative VIP-IN current-clamp recording in response to current injections (−10 to +20 pA, in 10 pA steps). Scale bars 20 mV and 250 ms. **(c)** Current-evoked action potentials (APs) were comparable in VIP-INs from female (♀) and male (♂) mice. **(d)** No difference in resting membrane potential (V_m_) between ♀ and ♂ mice. **(e)** Mouse sex did not affect VIP-IN rheobase. **(f)** VIP-IN membrane resistance (R_m_) was not different between ♀ and ♂ mice. **(g)** Hyperpolarization sag ratio was similar in VIP-INs from ♀ and ♂ mice. n/N = 23-27 cells from 5-7 mice per group.

To our knowledge, no published studies have systematically assessed whether VIP-INs display sex differences in physiology. We therefore performed a head-to-head comparison of several membrane properties following negative and positive current injections. Overall, we found no evidence for sex differences in membrane properties of PFC VIP-INs. VIP-INs from female and male mice displayed comparable current-evoked action potentials **(Figure 1c)**. VIP-IN resting membrane potential (V_m_) is comparable to other cell types within PFC, and no differences related to sex were detected **(Figure 1d)**. By contrast, relative to pyramidal cells and most other IN subtypes, VIP-INs displayed extremely low rheobase **(Figure 1e)** – i.e., the minimum current needed to elicit spike-firing. Additionally, VIP-INs exhibit extremely high membrane resistance (R_m_) **(Figure 1f)**, an overall measurement of excitability that is inversely proportional to the number of functional cell-surface ion channels. No sex differences in VIP-IN rheobase or membrane resistance were detected. Finally, we assessed hyperpolarization sag **(Figure 1g)**, a measurement of hyperpolarization- and cyclic nucleotide-gated (HCN) channels (Salling et al., 2018; Shah, 2014), which can regulate how certain subtypes of hippocampal INs respond to acute ethanol (Yan et al., 2009; Yan et al., 2010). Hyperpolarization sag was similar in VIP-INs from female and male mice. Taken together, these data indicate that VIP-INs are highly excitable cells without obvious basal sex differences in membrane physiology.

### 3.2 Acute ethanol application enhances excitability of PFC VIP-Ins

Previous studies in acute brain slices and organotypic cultures have shown that modest concentrations of ethanol can modify the physiology and function of neocortical and hippocampal INs (Carta et al., 2003; Harrison et al., 2017; Woodward and Pava, 2009; Yan et al., 2009; Yan et al., 2010). To probe whether ethanol directly can affect VIP-IN membrane physiology, we incubated acute PFC brain slices in 20 mM ethanol [∼0.09 BEC, comparable to peaks concentrations attained during IA ethanol (Salling et al., 2018)] for 1-5 hours before recording from VIP-INs **(Figure 2a)**. Comparable effects were observed in cells from female and male mice **(Figure S1)**, so data were pooled for the primary analysis. VIP-INs treated with ethanol fired more action potentials in response to positive current injections than ACSF control cells **(Figure 2b and 2C)**. Acute ethanol did not alter VIP-IN resting membrane potential **(Figure 2d)**; however, ethanol reduced rheobase **(Figure 2e)** and enhanced membrane resistance **(Figure 2f)** relative to matched control ACSF cells. Finally, hyperpolarization sag was not affected by ethanol application **(Figure 2g)**, suggesting that 20 mM ethanol does not acutely affect HCN channel function in VIP-INs in brain slices. Taken together, these findings suggest that ethanol can directly act on VIP-INs to enhance their excitability.

**Figure 2.**
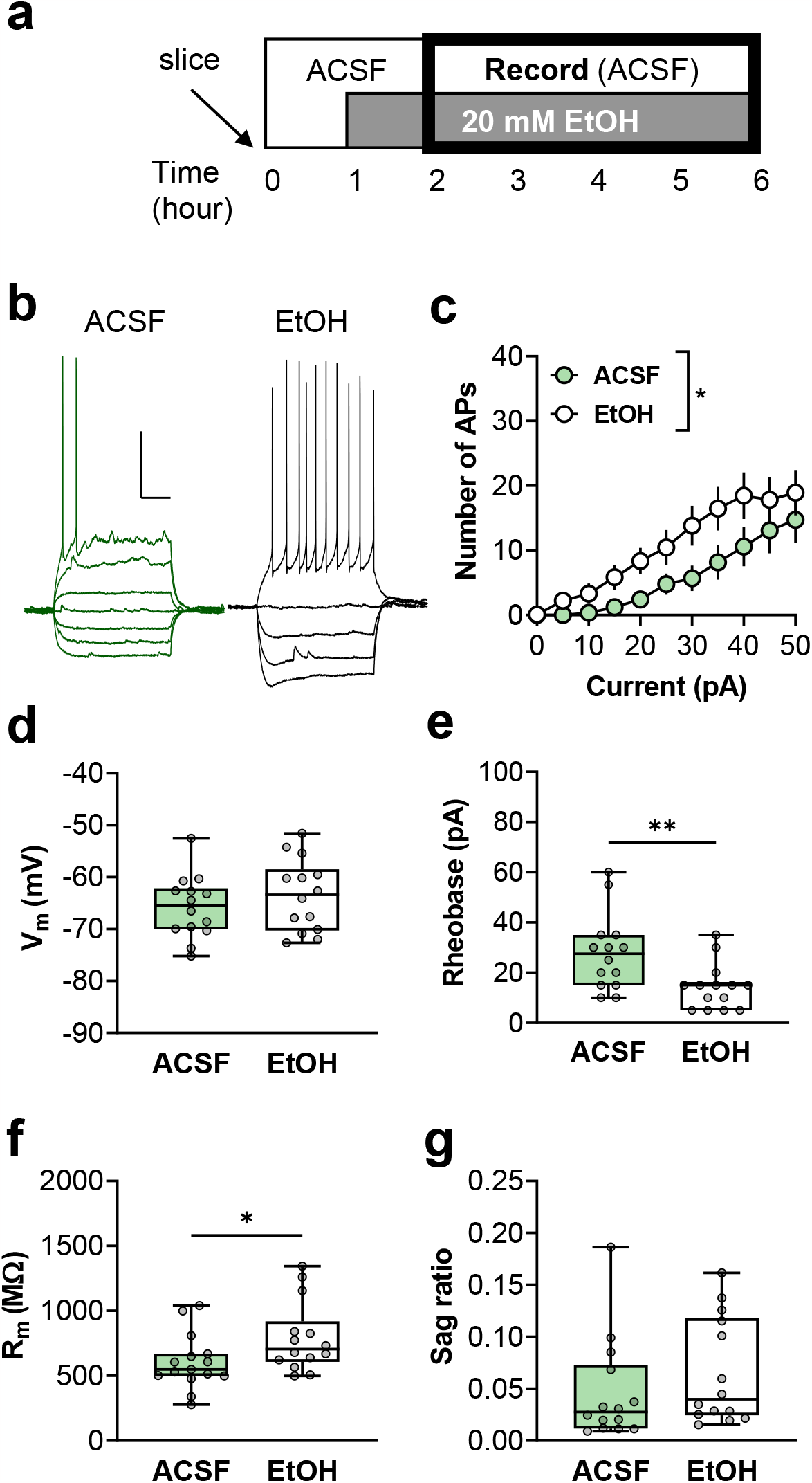
Acute ethanol application enhances excitability of PFC VIP-INs. **(a)** Experimental schematic. All slices recovered in ACSF for one hour, at which point half of the slices per mouse were transferred to ACSF supplemented with 20 mM ethanol. Recordings from ACSF or ethanol slices were then made 2-6 hours after sacrifice. **(b)** Representative VIP-IN current-clamp recordings from control (left, green) and 20 mM ethanol (right, black) slices. Traces depict responses to incremented positive current injections (−30 to +30/10 pA, in 10 pA steps). Scale bars 20 mV and 250 ms. **(c)** Acute ethanol enhanced VIP-IN current-evoked spiking (Two-way RM mixed-effects analysis: F_1,26_ = 6.5, *:p<0.05 main effect of IA ethanol). **(d)** VIP-IN V_m_ was not affected by acute ethanol. **(e)** Ethanol application decreased VIP-IN rheobase (Mann-Whitney test: U = 41.5, **:p<0.01). **(f)** Ethanol increased VIP-IN R_m_ (Mann-Whitney test: U = 44.5, *:p<0.05). **(g)** VIP-IN hyperpolarization sag was not affected by acute ethanol. n/N = 14 cells from 7 mice (4♀ / 3♂) per group.

### 3.3 Chronic voluntary drinking decreases excitability of PFC VIP-INs

The previous findings indicated that acute ethanol administration can alter VIP-IN excitability, so we next asked whether long-term drinking induces similar or related effects. We examined VIP-IN membrane physiology following intermittent access (IA) ethanol, a model that we and others have used to facilitate moderate-to-high levels of voluntary drinking in mice (Crabbe et al., 2012; Hwa et al., 2011; Joffe et al., 2020b, 2021; Salling et al., 2018). For alternating 24-hour periods, we provided VIP-tdTomato mice with free access to ethanol in their home cages **(Figure 3a)**. Female C57BL/6J mice generally drink more ethanol than male counterparts (Becker and Lopez, 2004; Finn et al., 2018; Joffe et al., 2020b; Jury et al., 2017). In the cohort used for these experiments, we observed slightly greater drinking in female mice, but the effect did not reach statistical significance **(Figure S2)**.

**Figure 3.**
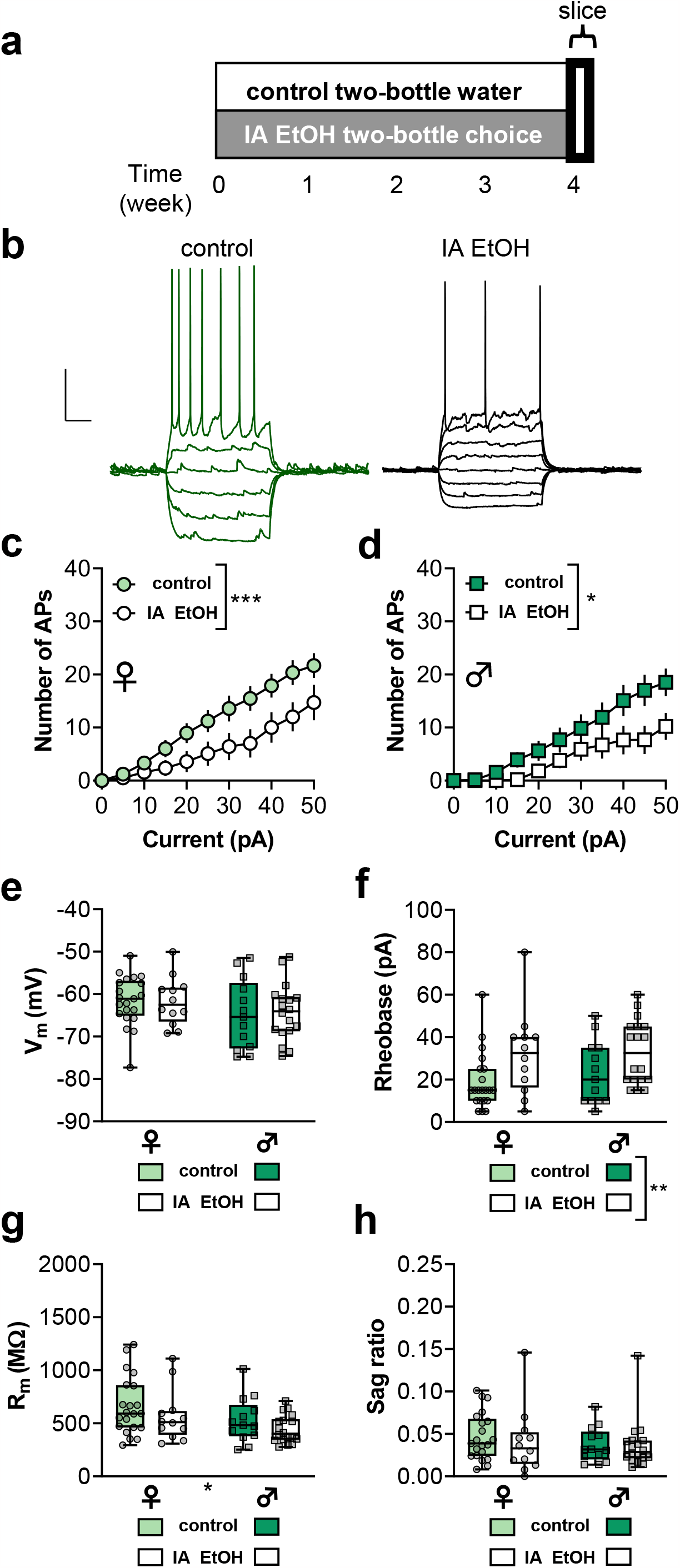
Chronic voluntary drinking decreases excitability of PFC VIP-INs. **(a)** Experimental schematic. VIP-tdTomato mice underwent 4 weeks of intermittent access (IA) ethanol two-bottle choice, or control two-bottle water, home cage drinking. Slices were prepared the day after the last drinking session. **(b)** Representative VIP-IN current-clamp recordings from control (left, green) and IA ethanol mice (right, black). Traces depict responses to incremented positive current injections (−30 to +20/40 pA, in 10 pA steps). Scale bars 20 mV and 250 ms. **(c/d)** IA ethanol decreased current-evoke spiking in VIP-INs in both ♀ mice (Two-way RM mixed-effects analysis: F_10,288_ = 3.2, ***:p<0.001 current x IA ethanol interaction) and ♂ mice (Two-way RM mixed-effects analysis: F_10,213_ = 2.3, *:p<0.05 current x IA ethanol interaction). **(e)** VIP-IN V_m_ was not affected by IA ethanol. **(f)** IA ethanol increased VIP-IN rheobase (Two-way ANOVA: F_1,60_ = 8.5, **:p<0.01 main effect of IA ethanol). **(g)** IA ethanol did not affect VIP-IN R_m_, but VIP R_m_ was lower in ♂ mice relative to ♀ mice in this cohort (Two-way ANOVA: F_1,60_ = 4.8, *:p<0.05 main effect of sex). **(h)** No effect of IA ethanol on hyperpolarization sag. n/N = 13-20 cells from 3-5 mice per group.

The day after the last drinking session, we sacrificed mice for brain slice electrophysiology and recorded from PFC VIP-INs. As with the acute ethanol studies, we examined VIP-IN membrane properties through a series of negative and positive current injections **(Figure 3b)**. While we previously found that acute ethanol enhances VIP-IN excitability, we observed the opposite effect following chronic drinking: VIP-INs from IA ethanol mice displayed decreased current-evoked spike-firing relative to matched cells from water-drinking controls **(Figure 3c and 3d)**. The resting membrane potential was not affected by IA ethanol or sex **(Figure 3e)**, suggesting that changes in spike-firing likely stem from a voltage-gated conductance. Consistent with the overall decrease in current-evoked spiking, drinking increased the minimum current required to elicit action potential firing in VIP-INs, as quantified by a greater rheobase in IA ethanol cells relative to controls **(Figure 3f)**. IA ethanol did not affect membrane resistance, although VIP-INs from male mice displayed reduced membrane resistance relative to cells from female mice in this cohort **(Figure 3g)**. Finally, no effect of drinking or sex was found on hyperpolarization sag **(Figure 3h)**. Taken together, these studies illustrate that chronic drinking generates opposing adaptations to VIP-INs relative to acute ethanol application.

### 3.4 Hypo-excitability of VIP-INs persists in male mice but not female mice

Several affective behavioral changes develop, and some behavioral changes wane, over the course of abstinence from voluntary drinking (Bloch et al., 2022; Holleran and Winder, 2017). We therefore aimed to understand whether drinking-associated changes to VIP-IN physiology persist into abstinence. To address this question, we provided a new cohort of mice with IA ethanol for 4 weeks, after which we removed all access to ethanol for 1-2 weeks before sacrificing animals for electrophysiology **(Figure 4a)**. In contrast to the studies performed without abstinence, we observed no differences in current-evoked spike-firing **(Figure 4b)**, or any other physiological parameter, in VIP-INs from female mice. VIP-INs from abstinent IA ethanol male mice, however, displayed decreased current-evoked spiking relative to control cells **(Figure 4c)**. Furthermore, VIP-INs from abstinent IA ethanol male mice had hyperpolarized resting membrane potentials relative to controls **(Figure 4d)**. Similar to previous studies at the early timepoint, VIP-INs from male mice in abstinence displayed greater rheobase **(Figure 4e)** and decreased membrane resistance **(Figure 4f)** than control cells. Finally, we assessed hyperpolarization sag potentials. VIP-INs from abstinent male mice displayed reduced hyperpolarization sag than controls **(Figure 4g)**, consistent with a potential reduction in HCN channel function. Overall, these findings suggest that VIP-INs from female mice may recover faster from the effects of chronic drinking when compared to male experimental counterparts. Furthermore, the changes in resting potential and hyperpolarization sag both suggest that additional adaptations in VIP-INs HCN channel function may develop during abstinence in male mice.

**Figure 4.**
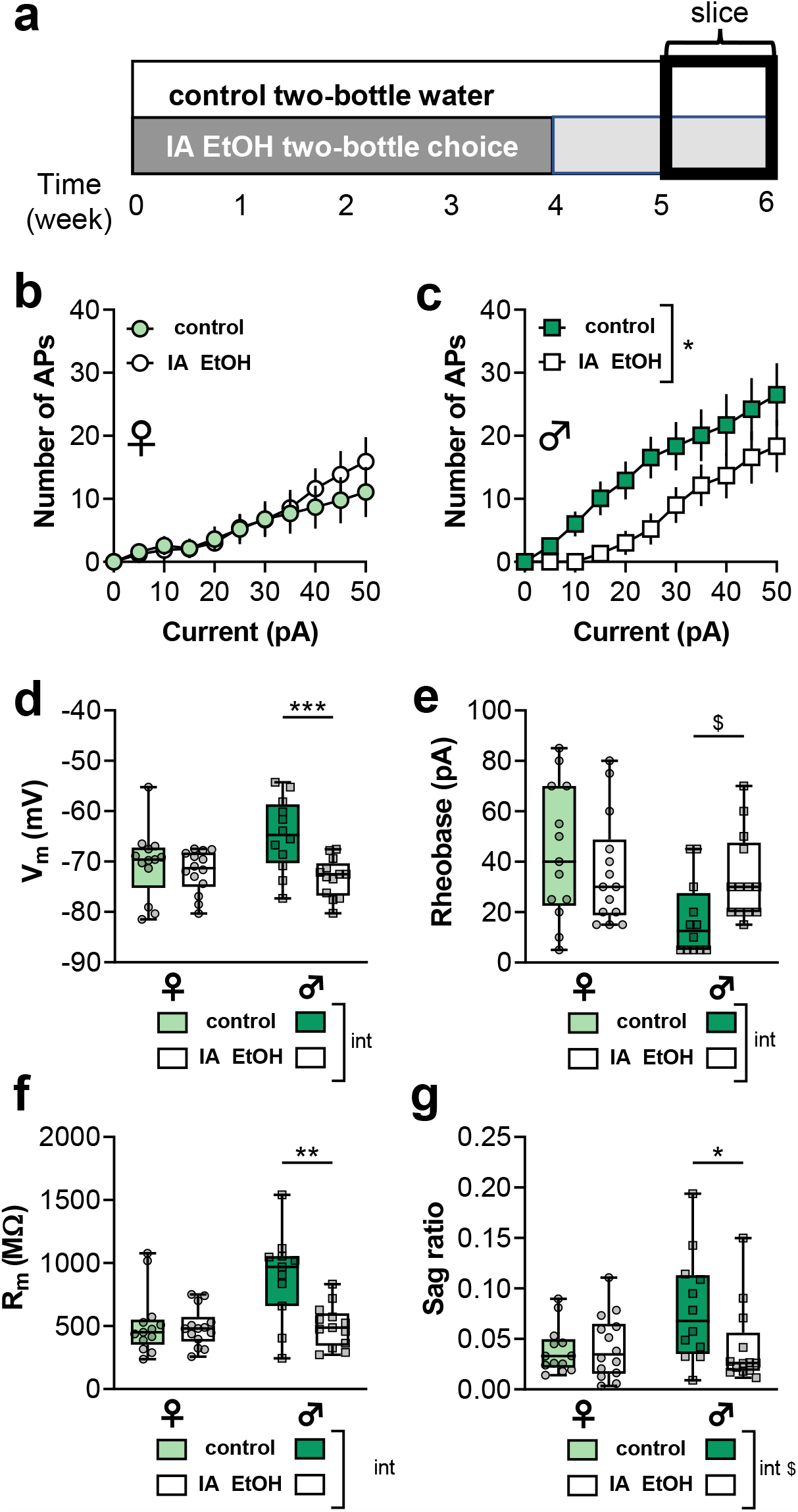
Hypo-excitability of VIP-INs persists in male mice but not female mice. **(a)** Mice underwent 4 weeks of IA ethanol in their home cages, followed by 1-2 weeks of abstinence. Animals were sacrificed for electrophysiology 7-11 days after the final drinking session. **(b)** No difference in current-evoked spiking in VIP-INs between control and abstinent IA ethanol ♀ mice. **(c)** VIP-INs from abstinent IA ethanol ♂ mice displayed reduced current-evoked spiking relative to water controls (Two-way RM mixed-effects analysis: F_10,228_ = 2.2, *:p<0.03 current x IA ethanol interaction). **(d)** In ♂ but not ♀ mice, VIP-INs displayed hyperpolarized V_m_ in abstinence from IA ethanol relative to water controls (Two-way ANOVA: F_1,48_ = 4.9, p<0.04 sex x IA ethanol interaction; ***: p<0.001 Bonferonni post test). **(e)** Abstinence from IA ethanol led to a trend increase in rheobase in VIP-INs from ♂ mice but not ♀ mice (Two-way ANOVA: F_1,48_ = 4.5, p<0.04 sex x IA ethanol interaction; $: p<0.1 Bonferonni post test). **(f)** In ♂ but not ♀ mice, VIP-INs displayed reduced R_m_ in abstinence from IA ethanol relative to water controls (Two-way ANOVA: F_1,47_ = 7.4, p<0.01 sex x IA ethanol interaction; ***: p<0.001 Bonferonni post test). **(g)** Abstinence from IA ethanol decreased hyperpolarization sag in VIP-INs from ♂ mice but not ♀ mice (Two-way ANOVA: F_1,48_ = 3.7, p<0.06 sex x IA ethanol interaction; *: p<0.05 Bonferonni post test). n/N = 12-14 cells from 3 mice per group.

### 3.5 Minimal effect of acute ethanol, IA ethanol, or abstinence on excitatory drive onto VIP-INs

In addition to intrinsic physiology, changes in synaptic strength can exert a major impact on a cell’s integration within its micro- and macro-neural circuits. Previous studies from our lab and others have described several changes to excitatory drive onto other IN subtypes in PFC (Dao et al., 2021; Ferranti et al., 2022; Joffe et al., 2020b; Trantham-Davidson et al., 2014; Trantham-Davidson et al., 2017). To examine excitatory drive onto VIP-INs, we electrically isolated sEPSCs by voltage-clamping cells near the reversal potential for GABA_A_ receptors. Acute ethanol did not affect the amplitude or frequency of sEPSCs in VIP-INs **(Figure 5a and 5b)**. Similarly, we found no effect of IA ethanol or sex on sEPSC parameters **(Figure 5c and 5d)**. After abstinence from IA ethanol, we observed a significant interaction between IA ethanol and sex on sEPSC amplitude, but post hoc comparisons were not significant **(Figure 5e)**. No effects of abstinence or sex were observed regarding sEPSC frequency **(Figure 5f)**. These data indicate that ethanol and chronic drinking do not have a major impact on excitatory synaptic properties on VIP-INs and suggest that the observed changes to membrane physiology occur in a cell-autonomous manner.

**5.**
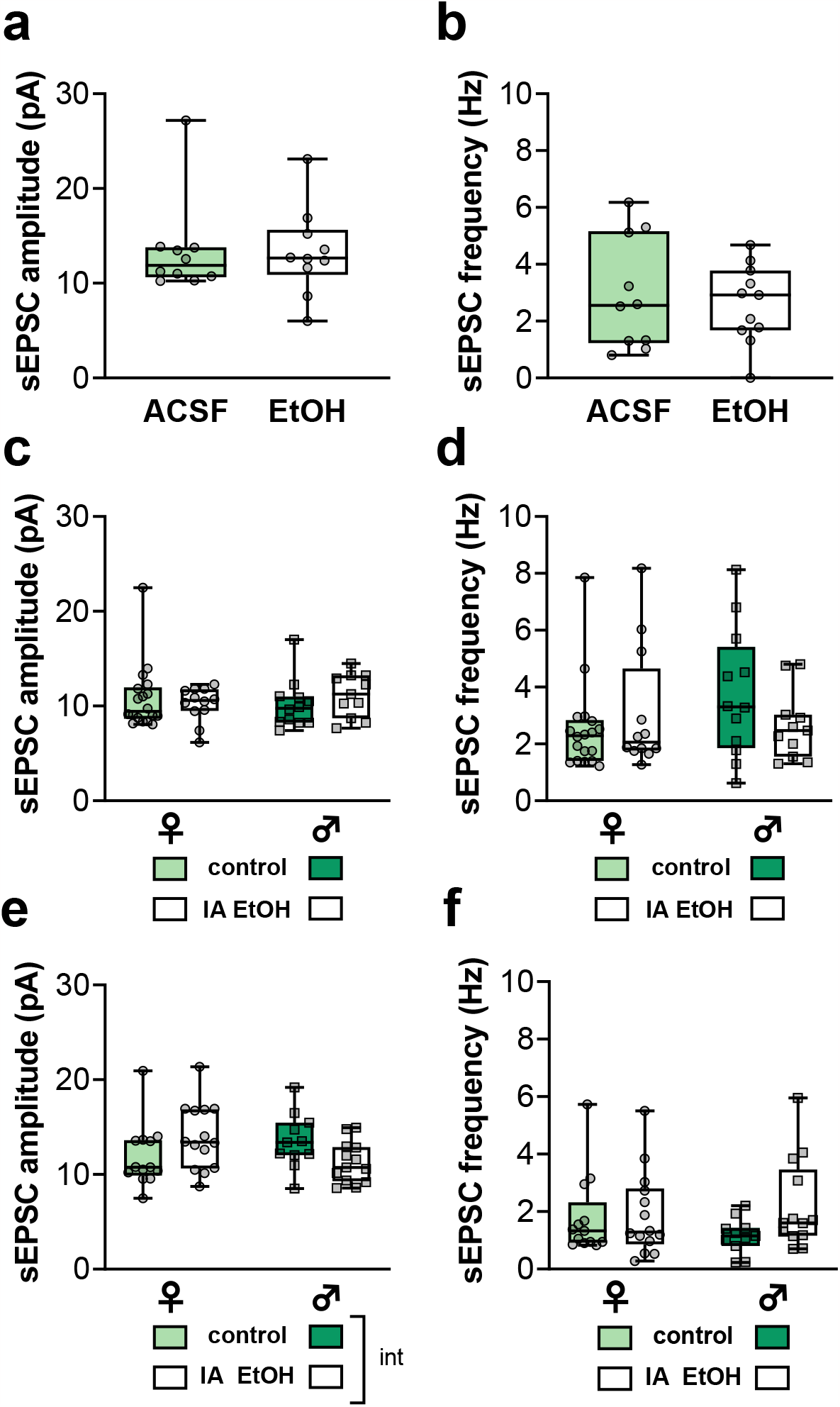
Minimal effect of acute ethanol, IA ethanol, or abstinence on excitatory drive onto VIP-INs. Spontaneous excitatory postsynaptic currents (sEPSCs) were measured in voltage-clamp configuration. **(a/b)** Application of acute ethanol (20 mM) did not affect the amplitude or frequency of sEPSCs on VIP-INs. n/N = 10 cells from 5 mice (3♀ / 2♂) per group. **(c/d)** VIP-IN sEPSC amplitude and frequency were comparable between IA ethanol mice and water controls. n/N = 13-20 cells from 3-5 mice per group. **(e)** An interaction between abstinence from IA ethanol and sex affected VIP-IN sEPSC amplitude (Two-way ANOVA: F_1,47_ = 6.4, p<0.02 sex x IA ethanol interaction), but post hoc comparisons did not reach significance (p>0.14). n/N = 12-14 cells from 3 mice per group. **(f)** No effect of abstinence from IA ethanol or sex on VIP-IN sEPSC frequency.

## 4 Discussion

The PFC is a key regulator of the acute and persistent effects of alcohol on brain function and behavior. PFC activity is dynamically modulated by VIP-INs, a unique type of neuron that inhibits other INs to enhance pyramidal cell activity (Audette et al., 2018; Lee et al., 2013; Pi et al., 2013; Williams and Holtmaat, 2019). To our knowledge, this series of studies represents the first published account of how ethanol and chronic drinking affect the function of PFC VIP-INs. Here, we found that ethanol acutely enhances the excitability of VIP-INs in brain slices. By contrast, following chronic voluntary drinking, VIP-INs displayed reduced spike-firing and greater rheobase relative to cells from water-only controls, consistent with a homeostatic reduction in excitability. Finally, we found that drinking-induced changes in VIP-INs persisted in male mice, but not female mice, over the course of one week of abstinence. Together, these studies identify unappreciated changes to PFC VIP-INs following acute ethanol and chronic drinking, paving the way for future studies to better understand the underlying mechanisms and to assess the relevance for problematic drinking behaviors.

Ethanol affects a wide variety of ion channels expressed in the central nervous system (Abrahao et al., 2017; Harrison et al., 2017). In these studies, we found that acute ethanol enhances the current-evoked spiking of PFC VIP-INs at a physiologically relevant concentration of 20 mM (∼0.09 BEC) **(Section 3.2)**. This effect could result from modulation of many targets, but we can begin to identify potential candidate molecules by evaluating the peripheral changes in membrane physiology. First, we observed that ethanol increased VIP-IN membrane resistance. As membrane resistance is inversely related to the number of functional ion channels on a neuronal membrane, it is likely that ethanol enhanced VIP-IN excitability by inhibiting one or more specific ion channels expressed by VIP-INs. Second, we found that ethanol decreased the rheobase and had minimal change on the resting membrane potential. These findings suggest that acute ethanol did not affect a tonic conductance, but instead acted on a voltage-gated channel. Taken together, this constellation of changes suggests that ethanol enhances VIP-IN activity by inhibiting a voltage-gated potassium current. Previous studies have found that ethanol inhibits KCNQ2 (or Kv7.2) currents at relevant concentrations (10-40 mM) in hippocampal pyramidal cells (Moore et al., 1990), in the ventral tegmental area (Koyama et al., 2007), and in heterologous expression systems (Cavaliere et al., 2012), supporting the hypothesis that KCNQ channels mediate ethanol’s excitatory actions on PFC VIP-INs. This hypothesis could have exciting ramifications for potential breakthrough treatments, as the KCNQ channel potentiator, retigabine, attenuates drinking in rodent studies (Knapp et al., 2014; McGuier et al., 2016), and PFC *Kcnq5* expression negatively correlates with voluntary drinking (Rinker et al., 2017). While their ability to modulate INs in PFC has not been documented, KCNQ channels regulate the timing and summation of spikes within hippocampal INs (Lawrence et al., 2006), providing more support for this untested hypothesis. BK channels represent another class of voltage-gated potassium channel that could be involved in the present effects on VIP-INs. While initial studies showed that ethanol enhances BK (or KCa1.1) channel function (Dopico et al., 1996), ethanol can also inhibit BK channels depending on post-translational modifications, intracellular calcium concentrations, and the co-expression of auxiliary subunits and associated proteins (Dopico et al., 2016).

While acute ethanol enhanced VIP-IN current-evoked spiking and decreased rheobase, we observed opposite effects following IA ethanol chronic drinking **(Section 3.3)**. These findings support the notion that drinking precipitates a homeostatic change in VIP-IN membrane physiology, potentially driven by the same ionotropic species(s) inhibited by acute ethanol. We also observed a sex-dependent change in VIP-IN excitability after abstinence: VIP-IN hypo-excitability persisted in male mice, but the membrane physiology of VIP-INs from abstinent IA ethanol male mice was similar to respective controls **(Section 3.4)**. One caveat to these findings is that the female control mice from the abstinence group appears to have reduced excitability relative to the female control groups in other studies. Based on this, it is possible that extended social isolation, or some other cohort-specific effect, may have prevented effects of ethanol from being detectable in this group of abstinent female mice. While these experiments were not designed to directly assess how group-versus single-housing might affect VIP-IN physiology, future studies should take this into account. Another interesting finding that emerged during abstinence is the hyperpolarization and reduced sag in VIP-INs from male mice. Both findings could be explained by a reduction in HCN channel function. Previous studies have found that IA ethanol during adolescence can induce membrane hyperpolarization and reduce sag currents in PFC pyramidal cells (Salling et al., 2018). While these effects have not been observed when IA ethanol is initiated in adulthood (Joffe et al., 2021; Salling et al., 2018), it is possible that VIP-INs may express unique HCN channel subunits and/or auxiliary proteins that render them sensitive to ethanol’s effects later in life. Support for this hypothesis can be found in previous studies in hippocampal INs. Ethanol has minimal effect on quiescent INs but increases spontaneous firing of active INs by facilitating HCN channel function (Yan et al., 2009; Yan et al., 2010). Thus, it is possible that chronic drinking may also induce a homeostatic change in HCN channel function in VIP-INs, at least in male mice. While we found no evidence for acute ethanol enhancing hyperpolarization sag or altering the resting potential in the present set of studies **(Section 3.2)**, high cell-to-cell variation of membrane physiology in VIP-INs and/or the limited sensitivity of experiments in current-clamp configuration may have precluded the ability to detect subtle effects on HCN channel function. Future studies might be designed to test whether ethanol modulates VIP-IN HCN channels by performing directed studies in voltage-clamp configuration and/or by pharmacologically blocking other conductances (e.g. potassium currents) that might otherwise obfuscate subtle effects. Finally, future studies should assess whether the long-term effects of ethanol on VIP-INs might involve ethanol’s ability to potentiate 5HT_3_ receptor function (Lovinger, 1999; Lovinger and White, 1991). From a translational perspective, investigating whether 5HT_3_ receptors are involved in ethanol’s actions on PFC interneurons in mice is critical, as cortical INs in humans do not express the 5HT_3_ receptor (Krienen et al., 2020). Overall, mechanistic studies describing how drinking reduces VIP-IN excitability will be an important area of future research towards the identification of potential novel treatment targets.

Previous studies from our laboratory and others have examined how ethanol exposure affects PFC IN subtypes derived from the medial ganglionic eminence (e.g., PV-INs and SST-INs) (Dao et al., 2021; Dao et al., 2020; Ferranti et al., 2022; Fish and Joffe, 2022; Hughes et al., 2020; Joffe et al., 2020b; Li et al., 2021; Trantham-Davidson et al., 2014; Trantham-Davidson et al., 2017). To our knowledge, however, this is the first report of ethanol’s effects on an IN subtype that arises from the caudal ganglionic eminence. These INs represent a broad and heterogenous group of neurons that preferentially reside in the superficial cortical layers (i.e., layers 1-3) (Krienen et al., 2020; Lee et al., 2010; Schuman et al., 2019; Tasic et al., 2018). Despite their importance in regulating cortical function, these cells have been tremendously understudied in biological psychiatry research, and there are many other subtypes to examine beyond VIP-INs. In fact, nearly 90% of INs in layer 1 do not express VIP (Schuman et al., 2019); examining how these IN subpopulations are affected by acute ethanol and chronic drinking represents a compelling direction for future research. Even VIP-INs themselves can be subdivided into further subtypes with important functional and molecular distinctions. At the highest level, VIP-INs can be dichotomized into generally non-overlapping subpopulations based on expression of cholecystokinin (CCK) or calretinin (CR) (Porter et al., 1998; Tasic et al., 2018). VIP/CR-INs are commonly bipolar cells that inhibit other interneurons (Fu et al., 2015; Lee et al., 2013; Melchitzky and Lewis, 2008; Pfeffer et al., 2013; Pi et al., 2013; Tasic et al., 2018). By contrast, VIP/CCK co-expressing INs are a type of basket cell that broadly dampens the activity of pyramidal cells through asynchronous GABA release (Freund et al., 1986; Kawaguchi and Kubota, 1998; Kubota and Kawaguchi, 1997). VIP/CCK-INs are more likely to reside in deeper cortical layers. Because the cells in the current studies all resided layers 1-3, the current results most likely relate to the function of disinhibitory VIP/ CR-INs. Nonetheless, considering their opposing effects on pyramidal cell activity, it would be worthwhile for future studies to compare effects on VIP/CR-INs vs VIP/CCK-INs. Finally, a significant subgroup of VIP-INs co-expresses choline acetyltransferase (ChAT) and the other requisite molecules for acetylcholine neurotransmission (Eckenstein and Baughman, 1984; Tasic et al., 2018). Recent functional studies have confirmed that VIP/ChAT-INs can directly excite other PFC cells via nicotinic receptors (Obermayer et al., 2019; von Engelhardt et al., 2007), thus they comprise an additional subpopulation that merits further study.

While significantly less is known relative to other IN subtypes, studies are beginning to describe how PFC VIP-INs regulate motivated and affective behaviors. The activity of VIP-INs is highly correlated with an animal’s state of arousal (Garcia-Junco-Clemente et al., 2017; Munoz et al., 2017), suggesting these cells may enhance the gain of salient information through cortical networks. Consistent with this idea, recent work has described how VIP-INs are recruited by reinforcing and punishing stimuli throughout all neocortical areas (Szadai et al., 2022). VIP-INs have been implicated in emotional behavior (Lee et al., 2019; Mossner et al., 2020), including aversiveness in response to neuropathic pain (Li et al., 2022). Based on this, it is tempting to speculate that changes in VIP-IN activity following IA ethanol are involved in affective disturbances and in heightened sensitivity to pain observed in many individuals with AUD. It will also be important to assess whether VIP-INs in other cortical areas respond to ethanol and adapt following chronic drinking. VIP-INs in somatosensory cortex regulate sensation, and these cells might be a substrate through which alcohol acutely alters vision, touch, and/or hearing. Therefore, future studies should certainly assess the functional relevance of alcohol’s effects on VIP-INs in multiple cortical areas.

## Funding and Declaration of Interest

This work was supported by the National Institutes of Health [grant number R00AA027806], the Whitehall Foundation [grant number 2022-08-005], and the Brain and Behavior Research Foundation. The authors declare no potential conflicts of interest.

## Acknowledgements

We thank members of the University of Pittsburgh Department of Psychiatry and Translational Neuroscience Program for stimulating discussions.

## Figure Legends

**S1.**
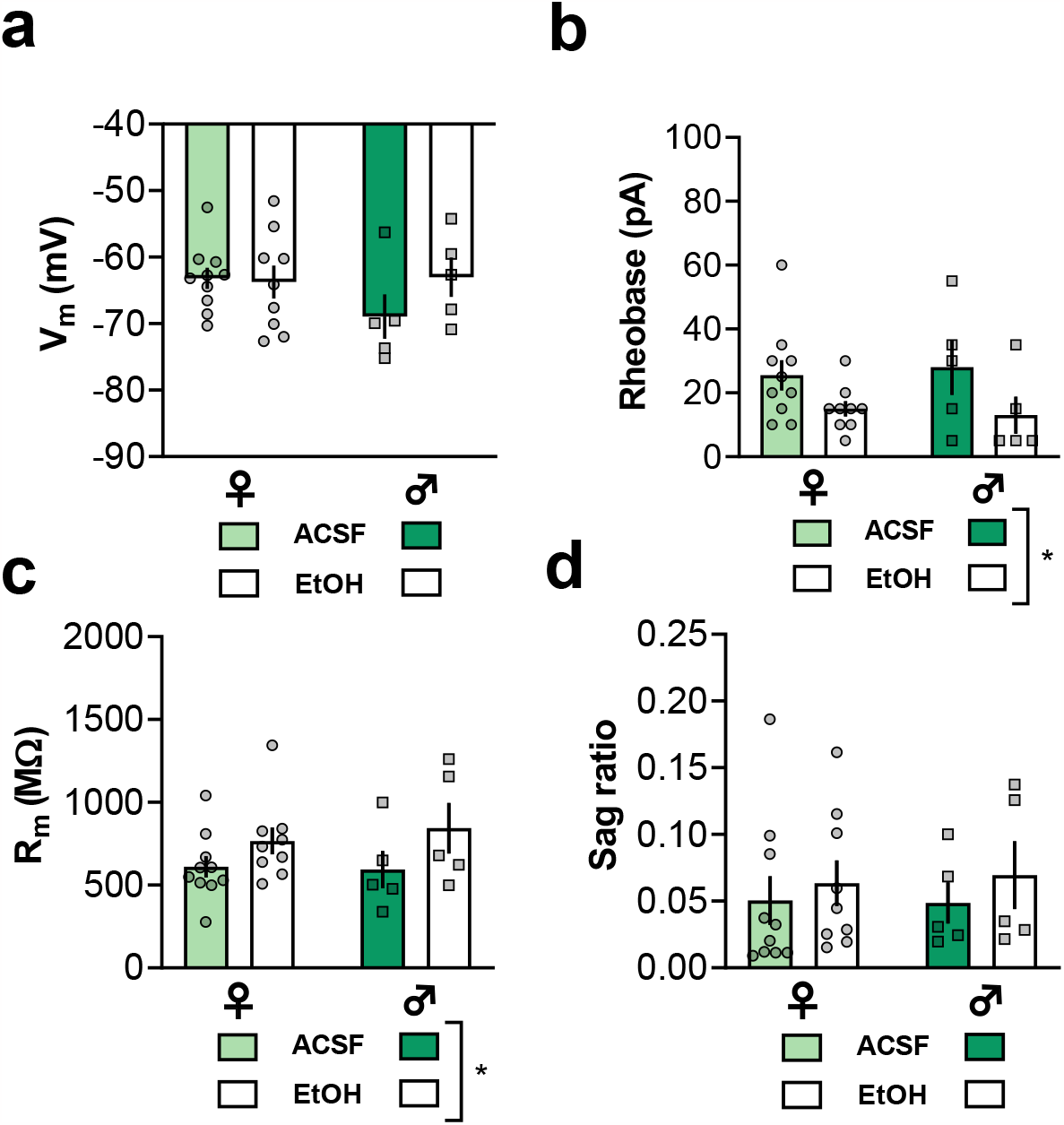
Related to Figure 2, No sex differences in actions of acute ethanol on VIP-IN membrane physiology. Data were pooled and presented in Figure 2. **(a)** No effect of sex or ethanol on VIP-IN V_m_. **(b)** Ethanol decreased VIP-IN rheobase comparably in ♀ and ♂ mice (Two-way ANOVA: F_1,25_ = 5.9, p<0.03 main effect of ethanol; F_1,25_ = 0.002, p>0.9 main effect of sex). **(c)** Ethanol increased VIP-IN R_m_ comparably in ♀ and ♂ mice (Two-way ANOVA: F_1,25_ = 4.3, p<0.05 main effect of ethanol; F_1,25_ = 0.09, p>0.7 main effect of sex). **(d)** No effect of sex or ethanol on VIP-IN V_m_.

**S2.**
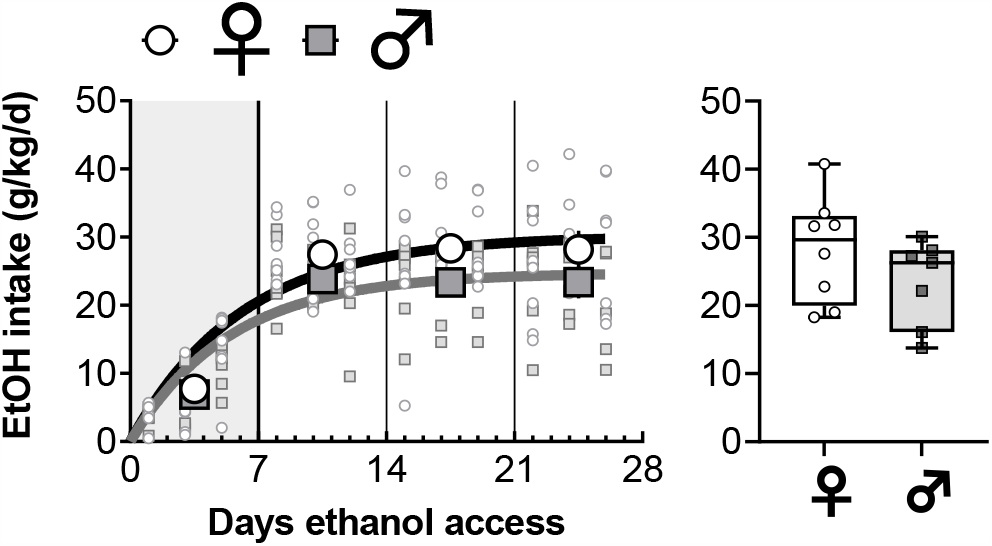
Related to Figures 3 and 4, Voluntary drinking for mice used in electrophysiology experiments. VIP-IN transgenic mice (backcrossed to C57BL6/J) began 4 weeks of two-bottle choice, voluntary drinking. Mice were separated by sex based on external genitalia. **Left**, one-phase association curves were fit to the ethanol intake data. The best-fit plateau for ♀ mice (27.8-33.2 g/kg) was greater than that for ♂ mice (22.8-27.0 g/kg) (F_1,176_ = 9.4, p<0.01). **Right**, the average intake over the fourth week of drinking was not different between sexes (t_13_ = 1.3, p>0.2). N = 7-8 mice per group.

## Notes

### Competing Interest Statement

The authors have declared no competing interest.

